# Expression of thioredoxin-1 in the ASJ neuron corresponds with and enhances intrinsic regenerative capacity under lesion conditioning in *C. elegans*

**DOI:** 10.1101/2022.09.19.508543

**Authors:** Noa W.F. Grooms, Michael Q. Fitzgerald, Binyamin Zuckerman, Samuel E. Ureña, Leor S. Weinberger, Samuel H. Chung

## Abstract

A conditioning lesion of the peripheral sensory axon triggers robust central axon regeneration in mammals. We trigger conditioned regeneration in the *C. elegans* ASJ neuron by laser surgery or genetic disruption of sensory pathways. Conditioning upregulates *trx-1* expression, as indicated by *trx-1* promoter-driven green fluorescent protein and fluorescence in situ hybridization, suggesting *trx-1* levels and associated fluorescence indicate regenerative capacity. Redox activity of *trx-1* functionally enhances conditioned regeneration, but both redox-dependent and –independent activity inhibit non-conditioned regeneration. Six strains isolated in a forward genetic screen for reduced fluorescence, which suggests diminished regenerative potential, also show reduced axon outgrowth. We demonstrate an association between *trx-1* expression and the conditioned state that we leverage to rapidly assess regenerative capacity.

## Introduction

The exquisitely complex and precise shape of neurons makes them highly susceptible to injury and disease. Compounding this weakness is the extremely limited regenerative capacity of the mammalian central nervous system (CNS), which severely limits functional recovery. Millions of people worldwide are affected by traumatic brain or spinal cord injury, amyotrophic lateral sclerosis (ALS), Alzheimer’s, Parkinson’s, or other neurodegenerative disorders (Feigin, *et al*., 2021; James, *et al*., 2019). The lack of effective treatments for these afflictions represents one of the areas of greatest need in modern medicine.

In a remarkable phenomenon known as lesion conditioning, damage to a neuron’s peripheral sensory fiber triggers cellular mechanisms to drive regeneration in the peripheral and central nervous systems (McQuarrie and Grafstein, 1973; Richardson and Issa, 1984). Lesion conditioning fosters a pro-regenerative environment (Dubový, *et al*., 2019; Xiong and Collins, 2012), enhances neuroregeneration (Hoffman, 2010) by overcoming CNS inhibitory growth cues (Chong, *et al*., 1996; Oudega, *et al*., 1994), and even dramatically reduces neurodegenerative markers in disease models (Franz, *et al*., 2009). Therefore, lesion conditioning drives and enables regeneration that could treat CNS afflictions.

Our laboratory utilizes the roundworm *Caenorhabditis elegans* for its strongly conserved genetics with mammals, facile genetics, optical transparency, and rapid development. In a prior study, we established a lesion conditioning model in the ASJ sensory neuron (Chung, *et al*., 2016). We discovered that mutation or pharmacological inhibition of genes in sensory transduction pathways can trigger conditioned regeneration, similar to the effect of a conditioning lesion. As shown in Tab. 1, these interventions produce several observable phenotypes, including conditioned regeneration and a form of ectopic axon outgrowth that has the same underlying genetic pathway.

**Table 1:** Interventions condition neurons by disrupting sensory signaling. Conditioning generates visible phenotypes. We examine *trx-1::gfp* fluorescence in this study.

| interventions → | condition<br>neuron → phenotype |
| --- | --- |
| surgery<br>conditioning mutations<br>pharmacology | regeneration<br>ectopic outgrowth<br>**fluorescence** |

In the prior study, we cell-specifically labelled the ASJ using a transgenic green fluorescent protein (GFP) reporter driven by a promoter of *trx-1*, an ortholog of human thioredo<u>x</u>in TXN. We noticed by eye that interventions that condition the ASJ also brighten the neuron, indicating an upregulation of the gene driving our fluorescent label. Our visual observations are consistent with prior studies. Following peripheral nerve axotomy, TXN accumulates in the nucleus and dramatically increases throughout the cytoplasm (Mansur, *et al*., 1998), indicating that it is a regeneration-associated gene (RAG) (Ma and Willis, 2015).

In this study, we quantify ASJ fluorescence and utilize it as an indicator of regenerative capacity. We quantitatively show that mutations that condition the ASJ by disrupting sensory signaling significantly brighten the neuron. We confirm this upregulation of *trx-1* by labeling mRNA transcripts with fluorescence *in situ* hybridization (FISH). Similarly, conditioning by lesion via laser surgery results in significantly increased fluorescence. The intensity of fluorescence following these interventions roughly corresponds with the neuron’s regenerative capacity. Additionally, we show that the *trx-1* gene restrains non-conditioned regeneration, under both redox-dependent and presumably redox-independent functions and enhances conditioned regeneration via redox-dependent functions. We leverage our fluorescent proxy to screen for mutations that underlie conditioning in the ASJ neuron. We isolate twelve mutant lines, spanning six putative mutations, with significantly reduced ASJ fluorescence. Six of these lines also display reduced ectopic outgrowth, indicating a disruption in the conditioning pathway. Our proxy enables efficient isolation of genes underlying lesion conditioning by enabling a visually based screen that does not require time-consuming surgery.

## Materials and methods

### *C. elegans* cultivation, strains, and mutagenesis

We followed established procedure for cultivation on agarose plates with OP50 bacteria at 20°C (Brenner, 1974). For fluorescent bead measurements, we cultivated animals on Bacto agarose plates at 20°C for multiple generations. We cultivated animals on Difco agarose plates at 20°C and 25°C for laser surgeries and ectopic outgrowth experiments, respectively. We confirmed the genotypes for all mutant strains by polymerase chain reaction followed by Sanger sequencing through GENEWIZ (for single-nucleotide polymorphisms) or gel electrophoresis (for large deletions). Tab. 2 lists the genetics of the strains utilized. We followed established protocol for ethylmethanesulfonate (EMS) mutagenesis (Brenner, 1974).

**Table 2:** Strains and genotypes utilized.

| mutation | translational marker |  | transcriptional marker |
| --- | --- | --- | --- |
|  | *ofls1[ptrx-1::trx-1::gfp]* | *ofls4[ptrx-1::trx-1::gfp]* | *ofls5[ptrx-1::gfp]* |
| none | BOS2 | OE3417 | BOS41 |
| *dlk-1(ju476)I* | BOS71 |  | BOS44 |
| *dlk-1(ju476)I; trx-1(ok1449)II* |  |  | BOS42 |
| *dlk-1(ju476)I; trx-1(syb206)II* |  |  | BOS82 |
| *egl-19(n582)IV* |  | BOS70 |  |
| *sax-1(ky211)X* | BOS74 |  |  |
| *tax-2(p691)I* | BOS72 | BOS76 |  |
| *tax-4(p678)III* |  | BOS73 |  |
| *trx-1(ok1449)II* |  |  | VZ763 |
| *trx-1(syb206)II* |  |  | BOS80 |
| *unc-36(e251)III* | BOS69 |  |  |
| *unc-43(n1186n498)IV* |  | BOS75 |  |

### FISH labeling and imaging

We used the designer tool from Stellaris (LGC Biosearch Technologies) to create probes (Table S1) that detect *trx-1* transcripts. We designed the probes under the following settings: a masking level of 5 (to avoid cross-hybridization to commonly expressed RNAs at high levels) and at least 2 base pair spacing between single probes. We generated 16 probes (CAL Fluor® Red 590 conjugated) for mature *trx-1*, targeting all exons and both UTRs to allow maximal probe number. Of note, we were unable to reach the optimal amount of probes recommended by the manufacturer (>24) due to the short mature length of *trx-1*. We synchronized animals by bleaching, hatched overnight in M9 buffer, and grown until young adult stage in standard conditions on NGM plates. We used fixation and hybridization protocol as described (Bolkova and Lanctot, 2016). We mounted animals mounted with ProLong™ Glass Antifade Mountant with NucBlue™ Stain (P36983, Invitrogen™). We acquired images using an Evident Scientific (formerly Olympus) FLUOVIEW FV3000 laser scanning confocal microscope, equipped with a high-sensitivity spectral detector unit (with Peltier-cooled GaAsP photomultiplier modules) and X Line UPLXAPO60XO oil objective. For each animal, we acquired 17 z-planes with a step size of 0.4 μm.

### Laser surgery, imaging of axon regeneration and ectopic outgrowth, and postprocessing

We followed established procedures for animal immobilization by sodium azide (Chung, *et al*., 2006) and femtosecond laser surgery (Harreguy, *et al*., 2020; Wang, *et al*., 2022). We severed only the axon (within 2 μm from the cell body), only the dendrite (2/3^rd^ the length distal from the cell body), or both the axon and dendrite (axon+dendrite), and we reimaged 48 hours afterwards. We followed established procedures for immobilizing animals on agarose-azide pads, imaging axon regeneration and outgrowth, and postprocessing images to measure regenerated length (Chung, *et al*., 2016). Note that neurons in certain genetic backgrounds may appear dim in some images due to the uniform normalization needed for comparison between backgrounds. These neurons are sufficiently bright to unambiguously measure the regenerated length, particularly after laser surgery, which upregulates *trx-1*-driven GFP.

### Imaging with fluorescent beads

Fluorescence measurements are affected by numerous factors, including the intensity of the light source and transmission through microscope optics. These factors may change over time as the components age and their alignment degrades. To account for these effects, we calibrated fluorescence measurements by imaging animals alongside fluorescent beads and normalizing measurements across the images taken. We thoroughly vortexed 0.3% and 3% 2.5 μm stock bead solution (Invitrogen I7219) and mixed it with nematode growth medium (NGM) buffer (Chung, *et al*., 2016) in a 5:1 NGM to bead volumetric ratio. We dropped 5.0 μL of the 5:1 solution to the center of the agarose-azide pad, and then picked adults to the bead solution droplet. To approximately match neuron to bead fluorescence, we used 3% beads for *tax-2* and *tax-4* strains and 0.3% beads for other strains and surgical conditions. We imaged worms through a Nikon 0.30 NA, 10x microscope objective and an Andor Zyla sCMOS Plus camera. Z-stack images were captured with 1-µm spacing.

### Postprocessing of fluorescence measurements

We create maximum intensity projections of our raw z-stack images in ImageJ and identify the brightest pixel within the ASJ cell body and dendrite for each animal. The pixel values are the raw fluorescent intensities. Additionally, we average the brightest pixel value within five representative beads in each raw image. We normalize and scale all raw neuron intensities to our reference (*tax-2*) intensities according to equation (1):

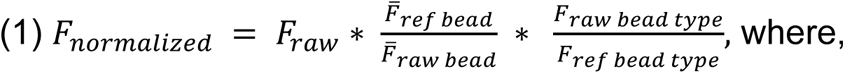

*F_raw_* is raw fluorescent intensity of ASJ cell body or dendrite.

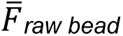 is the average of the fluorescence intensity of five representative beads in the raw image.

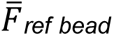 is average of the fluorescence intensity of all representative beads in the set of reference images.

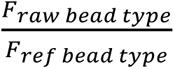 is the true ratio of bead fluorescent intensities according to manufacturer data. This ratio is 0.08 if the raw image is calibrated with 0.3% beads or 1.00 if calibrated with 3% beads like the reference image.

The first ratio of Eq. (1) normalizes for day-to-day variability in imaging parameters between an image and the reference image set. If beads appear unusually bright in an image due to stronger illumination or increased transmission, then this ratio will compensate. If the beads are abnormally dim, the opposite will occur. The second ratio compares the expected brightness of the bead type used in the image against the reference bead type. Under ideal imaging conditions, this second ratio comparing the bead intensities from manufacturer data should be the inverse of the first ratio that compares the measured bead intensities. Therefore, the product of the two ratios should center around 1.0 if the bead intensities match their manufacturer specifications. To quantify FISH signal, we adapted an established procedure (Iourov, *et al*., 2005).

We measured the intensity of the hybridization fluorescence in ImageJ by creating maximum intensity projections of our z-stacks and acquiring the brightest pixel value for each ASJ cell body. Since *tax-2* strongly upregulates *trx-1*, we could not quantify the signal at a single molecule (smFISH) level.

### Statistics and interpretation of results

For axon length and fluorescence measurements, we calculated *P* values by the unpaired, unequal variance, two-tailed *t* test. The conditioning effect is the difference between regenerated length after axon cut and after axon+dendrite cut. Its standard deviation is the square root of the sum of the squares of the regenerated length standard deviations. To compare conditioning effects in different backgrounds, we calculated *P* values by the unpaired, unequal variance, two-tailed *t* test. To compare frequencies of ectopic outgrowth, we used Fisher’s exact test.

## Results

### Defects in sensory transduction upregulate *trx-1* expression in the ASJ neuron

The ASJ is an amphid sensory neuron located in the nose of *C. elegans* (Fig. 1a, left). The amphids are bilateral, bipolar neurons following a stereotyped morphology consisting of a cell body, a sensory dendrite that extends to the nose, and an axon that mediates synaptic connections in the nerve ring (White, *et al*., 1986). The ASJ axon regenerates when it is severed. Mutation of dual-leucine kinase, *dlk-1*, completely abolishes what we call “conventional” (*i.e.*, single-axotomy) axon regeneration (Chung, *et al*., 2016). The regeneration of other *C. elegans* (Ghosh-Roy, *et al*., 2010; Hammarlund, *et al*., 2009) and mammalian neurons (Itoh, *et al*., 2009; Shin, *et al*., 2012) also show a strong dependence on *dlk-1*. We trigger DLK-independent regeneration by concomitantly severing the ASJ axon and dendrite in our lesion conditioning model. This DLK-independent regeneration exhibits several hallmarks of lesion-conditioned regeneration, including mediation by cyclic adenosine monophosphate (cAMP) (Neumann, *et al*., 2002; Qiu, *et al*., 2002) and inhibition by L-type voltage gated calcium channels (Enes, *et al*., 2010). We also uncovered several sensory mutations that significantly reduce ASJ neuronal activity levels and thus condition the ASJ to regenerate in *dlk-1* mutants without a physical lesion (Chung, *et al*., 2016).

**Figure 1:**
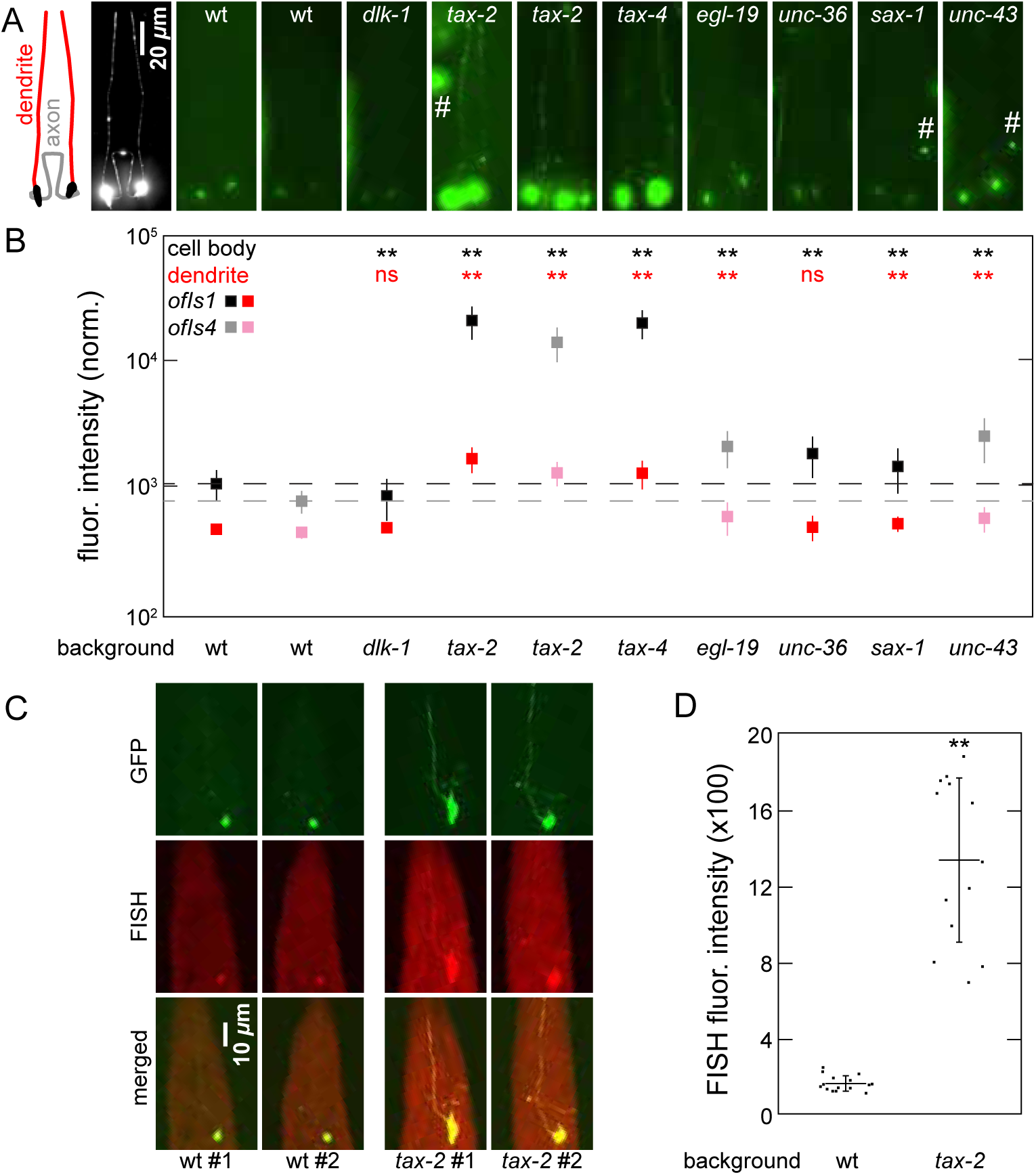
Sensory mutations upregulate ASJ neuron expression of *trx-1*. (a) Leftmost panels: Line drawing and typical ASJ neuron. Remainder of panels: representative images of *ptrx-1::trx-1::gfp* in wild-type (wt), mutant backgrounds. Sensory mutations, particularly *tax-2* and *tax-4*, upregulate expression. Identical brightness, contrast for all images except leftmost. Fluorescent beads indicated by #. (b) Quantification of fluorescence in part A. Data represented as average ± standard deviation (SD). *n =* 50 for each data point. ** *p < 0.001* (c) FISH confirms upregulation of *trx-1* in sensory mutant. Representative images of *trx-1*-targeted labels in GFP (top), FISH (middle), and merged (bottom) in wild-type and *tax-2* animals. (d) Quantification of fluorescence in part C. Each point represents measurement in one neuron. Data represented as average ± SD.

We characterized the effect of a subset of these mutations on the expression of *ptrx-1::trx-1::gfp* translational fusion reporters *ofIs1* and *ofIs4*. The loci of crucial genes *egl-19* and *unc-43* are on chromosome IV close to the integration site of *ofIs1*, which necessitated the use of *ofIs4*. While our previous study utilized an extrachromosomal array reporter, *ofIs1* and *ofIs4* are integrated into the genome, allowing more accurate comparison of expression levels across animals. We generated fluorescent animals with the mutations by crossing in *ofIs1* and *ofIs4* (Fig. 1a) and measured brightness of the ASJ cell bodies and dendrites to roughly assess TRX-1 levels and localization at a subcellular level (Fig. 1b). Scattered and out-of-focus light from the larger, brighter cell body often hides fine or dim structures nearby. Thus, we were unable to consistently measure the intensity of the axon. In wild-type (wt) animals, the ASJ cell body and dendrite are relatively dim. In a *dlk-1* mutant background, the ASJ fluorescence remains dim with a slight decrease in cell body fluorescence (*p < 0.001*), consistent with a non-conditioned regenerative state. The remainder of the mutations significantly increase the brightness of the ASJ fibers and especially cell bodies (*p < 0.001* for all mutations). The most dramatic alteration in fluorescent intensity occurs under *tax-2* and *tax-4* (Fig. 1b, *p < 0.001*), which also produce the strongest conditioned regeneration of the sensory mutations we studied. The fluorescence in the dendrites of each conditioned strain significantly increase except for *unc-36*, which may be consistent with the weaker conditioning effect of *unc-36* (Chung, *et al*., 2016). We compare *trx-1* expression via fluorescence with regenerative capacity in several conditioned backgrounds (Fig. S1). For ASJ cell bodies and dendrites, there is a general positive correlation between regeneration and *trx-1* expression. However, the correlation is not strong enough to reliably predict each mutation’s impact on regeneration. Thus, the expression of *ptrx-1::trx-1::gfp* in the ASJ generally corresponds with regenerative potential but is not predictive at a single gene level.

We confirmed the upregulation of *trx-1* by targeting *Trx-1* mRNA with FISH probes. In *C. elegans*, *trx-1* is significantly expressed in only the ASJ neuron and the intestines. We labeled transcripts in wild-type and *tax-2*, our strongest conditioning mutation, animals. As shown in Fig. 1c, to verify our FISH probes accurately targeted *trx-1*, we imaged the ASJ GFP in green (top row) and FISH in red (middle row). The overlays demonstrate clear colocalization of signals in 3D, indicating the probes accurately target *trx-1* (bottom row). We quantified fluorescence of the FISH hybridization following the procedure used for GFP images in Fig. 1a. We measured an 8x enhancement in ASJ fluorescence in *tax-2* animals compared to wild-type (Fig. 1d). This increase is similar to what we see in *ofIs1* and *ofIs4* fluorescence, further confirming the upregulation of *trx-1* in response to conditioning.

### Dendrite cuts upregulate *trx-1* expression and stimulate *dlk*-independent regeneration in the ASJ neuron

We also examined changes in ASJ fluorescence in *dlk-1* mutant two days after cutting the axon, the dendrite, or the axon and dendrite concomitantly (denoted as “axon+dendrite” or “a+d”). Figure 2a shows images of the ASJ in *dlk-1* animals 48 hours after surgery as well as in a chronically conditioned background, *tax-2*, to represent a strongly conditioned fluorescence level. Figure 2b shows the measured cell body and dendrite fluorescent intensities after each surgery. Our imaging procedure without surgery (mock condition) does not significantly alter ASJ fluorescence. Surgical interventions on the axon, dendrite, or axon+dendrite significantly increase fluorescence. Notably, severing the dendrite or axon+dendrite significantly increases ASJ fluorescence (*p < 0.001*) of nearly the entire population to a brightness that begins to approach the level in *tax-2* animals (right side, Fig. 2b). The fluorescence after axon+dendrite surgery is not significantly different from after dendrite surgery (*p = 0.163*), suggesting that cutting the dendrite is sufficient for activating the conditioning response. Figure 2c scatter plots the dendrite fluorescence against the cell body fluorescence post-surgery, clustering neurons by fluorescent intensity for comparing to conditioned state. Data points tightly clustered on the left side of the plot (dimmer ASJ fluorescence, mock and axon cut) represent a non-conditioned state. Interestingly, changes in fluorescence after axon cut exhibit bimodal distribution where half remain as dim as the ASJ in mock animals and the other half noticeably brighten. The cell body and the dendrite fluorescent intensities are correlated, leading to a clear demarcation between non-conditioned and conditioned states. Figure 2c confirms that neurons are strongly brightened following dendrite and axon+dendrite surgeries. The heightened levels of fluorescent intensity following dendrite or axon+dendrite cut are consistent with these interventions’ abilities to condition the neuron to regenerate. These results are consistent with published findings that TXN upregulates following peripheral axon injury, the hallmark of a RAG (Bai, *et al*., 2003; Mansur, *et al*., 1998).

**Figure 2:**
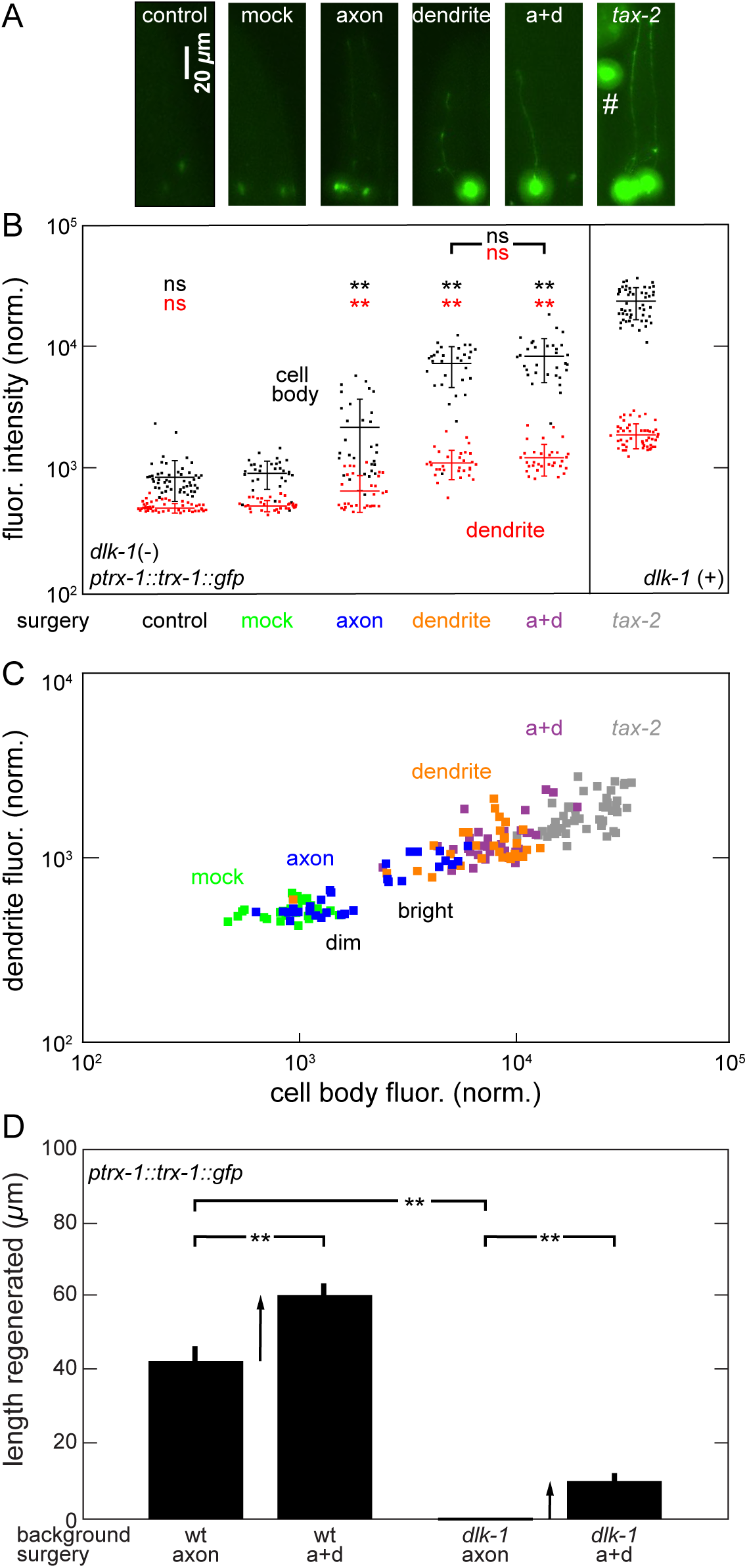
Dendrite cuts upregulate *trx-1* expression and condition ASJ to regenerate. (a) GFP fluorescence images of ASJ neuron. Fluorescent bead indicated by #. Identical brightness, contrast for all images. (b) Quantification of fluorescence in part A. Data represented as average ± SD. Each dot represents one animal. (c) ASJ cell body and dendrite fluorescence distributions from part B. Dendrite cut brightens fluorescence in entire population. (d) Total length of ASJ regeneration following axon or axon+dendrite (a+d) cut in wt, *dlk-1*. Dendrite cut enhances ASJ regeneration under DLK-independent mechanism. Arrows represent conditioning effect, or contribution of conditioning to additional regenerated length from dendrite cut. Data represented as average ± standard error of the mean (SEM). *n* ≥ 20 for all conditions. ** *p < 0.001*. Control and *tax-2* images and data replicated from Fig. 1.

Postsurgery regenerated length in ASJ expressing the translational reporter *ofIs1* match regeneration under a different translational reporter (Chung, *et al*., 2016). Conventional regeneration in the ASJ is largely *dlk*-dependent. Figure 2d displays length regenerated by the ASJ in wild-type and *dlk-1* following axotomy or axon+dendrite cut. The ASJ regenerates 43 μm on average following axotomy in wild-type but does not regenerate in *dlk-1* (*p < 0.001*). The conditioning effect is fundamentally the increase in regeneration arising from a dendrite cut, indicated by the arrows in the figure. Under our model, axon+dendrite cut triggers conventional and conditioned regeneration, which both contribute to the total length. In both wild-type and *dlk-1* background, the addition of the dendrite significantly increases regenerated length (*p* < 0.001) indicating a significant conditioning effect. Because *dlk-1* eliminates conventional ASJ regeneration, it exposes regeneration due to conditioning. Thus, we associate regeneration in *dlk-1* with conditioned regeneration. Note also that while axotomy alone does not produce conditioned regeneration in *dlk-1*, it significantly increases fluorescence of some neurons. This divergence indicates that the fluorescent reporter does not fully indicate the conditioned regenerative state and that *trx-1* expression alone is not sufficient for activating conditioned regeneration. We speculate that the significant increase in fluorescence after axotomy alone could be an upregulation of *trx-1* in response to neuroinjury without activating conditioning mechanisms.

### Opposite effects of *trx-1* on two forms of ASJ regeneration

Given that TXN is a RAG (upregulated following neuron injury), we wanted to determine if *trx-1* is involved in either form of regeneration. Our prior work utilized a translational *ptrx-1::trx-1::gfp* fusion marker (*ofIs1 or ofIs4*), potentially resulting in an overexpression of functional TRX-1 protein in the ASJ. Therefore, we visualized the ASJ with a transcriptional marker *ofIs5[ptrx-1::gfp]* and probed the effects of *trx-1(ok1449)*, which is a loss-of-function allele deleting a 860 bp region spanning part of the proximal promoter region up to the last exon (Miranda-Vizuete, *et al*., 2006). As shown in Fig. 3, the trends in regeneration in wild-type and *dlk-1* are consistent with regeneration under translational fusion reporters in Fig. 2 and in our prior study (Chung, *et al*., 2016). In wild-type animals, the axon regenerates at a basal rate when it is severed, and mutation of *dlk-1* eliminates conventional regeneration. The addition of a dendrite cut increases regeneration in wild-type (*p < 0.05*) and restores regeneration in *dlk-1* due to the conditioning effect (*p < 0.001*).

**Figure 3:**
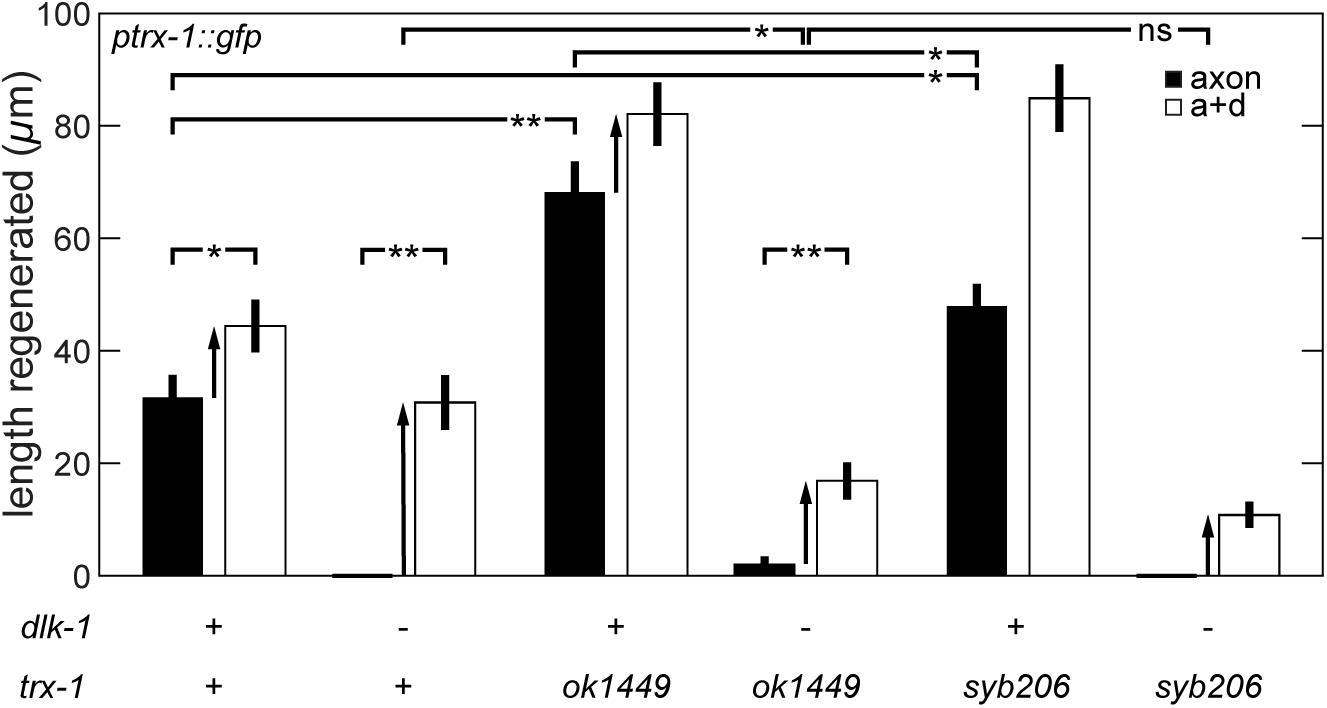
*trx-1* modulates ASJ regeneration. Mutation of *dlk-1* and *trx-1* differentially modulate regeneration after axon or a+d surgery. ok1449 is *trx-1* null allele; syb206 is *trx-1* redox-dead allele. Arrows represent conditioning effect. Redox-dependent and – independent functions of *trx-1* modulate conventional and conditioned regeneration. Data represented as average ± SEM*. n ≥ 20* for all conditions. * *p < 0.05, ** p < 0.001*.

ASJ conventional regeneration in *trx-1(ok1449)* is significantly stronger than in wild-type (*p < 0.001*), suggesting that *trx-1* restrains conventional regeneration. Introducing the *dlk-1* mutation to *trx-1* mutant background nearly eliminates conventional regeneration (Fig. 3), similar to results in translational fusions and *trx-1(+)*. The conditioning effect in *dlk-1; trx-1(ok1449)* is significantly less than the conditioning effect in *dlk-1*, as indicated by the arrows in Fig. 3 (*p < 0.05*). These data suggest that *trx-1* enhances conditioned regeneration.

### Redox functions of *trx-1* modulate both pathways of ASJ regeneration

To further uncover the role of *trx-1* in governing regeneration in the ASJ, we examined its redox-dependent functions. Two cysteine residues at conserved active sites on thioredoxin enable reversible oxidation (Lee, *et al*., 2013). Substituting one or both cysteine residues with serines, a redox-inactive amino acid, abolishes thioredoxin’s redox activity. In *C. elegans*, *trx-1(syb206)* mutants with both residues substituted (wild-type CGPC → redox-inactive SGPS) were utilized to demonstrate that *trx-1* regulates development through separate redox-dependent and –independent pathways (Sanzo-Machuca, *et al*., 2019). We assessed regeneration in the redox-inactive mutant to isolate thioredoxin’s redox-dependent effects (Fig. 3). Following axon cut only, regenerated length in *trx-1(syb206)* is significantly longer than in wildtype (*p < 0.05*) but significantly shorter than in the full *trx-1(ok1449)* null (*p < 0.05*). These results suggest that conventional regeneration is inhibited by *trx-1* under both redox-dependent and putatively redox-independent functions. In contrast, there is no significant change in the conditioning effect (indicated by arrows) between *dlk-1; trx(ok1449)* and *dlk-1; trx-1(syb206)* (*p = 0.141*). This result suggest that conditioned regeneration is likely mediated solely by redox-dependent functions of *trx-1*.

### Screening of *trx-1* reporter isolates mutants with reduced regenerative potential

One powerful technique for exposing genes underlying a phenotype is the unbiased forward screen (Brenner, 1974). In brief, a screen stochastically introduces mutations into the genomes of many animals. The correlation between *trx-1* expression and regenerative capacity suggests that we can rapidly screen for genes involved in conditioned regeneration utilizing a *trx-1* fluorescent reporter. We exposed our chronically conditioned *tax-2* strain, labeled with the translational reporter *ofIs1*, to ethyl methanesulfonate (EMS). Starting in the F_2_ generation, we isolated mutant lines by visually screening for decreased fluorescence in the ASJ neuron by eye (Fig. 4a) which potentially indicates a defect in a conditioning-related gene. Often, it was difficult to detect a significant reduction in cell body fluorescence, so we screened for significantly dimmer dendrites that we could not observe at a set magnification. We generated twelve mutant lines, spanning six putative mutations, with significantly reduced ASJ fluorescence in the cell body (Fig. 4b) or dendrite (Fig. 4c). In a typical screen, there is a frequency of roughly one null mutation in a given gene for every 2000 F_1_s screened (Jorgensen and Mango, 2002). Given that we screened 100 F_1_s, it is highly unlikely (though possible) that our screen produced the same mutations or mutations in the same gene. Thus, by the statistics, we strongly expect the six F_1_ lines isolated to carry distinct mutations in distinct genes. Ten of these lines display significantly reduced fluorescence in both the dendrite and cell body (*p < 0.001*).

**Figure 4:**
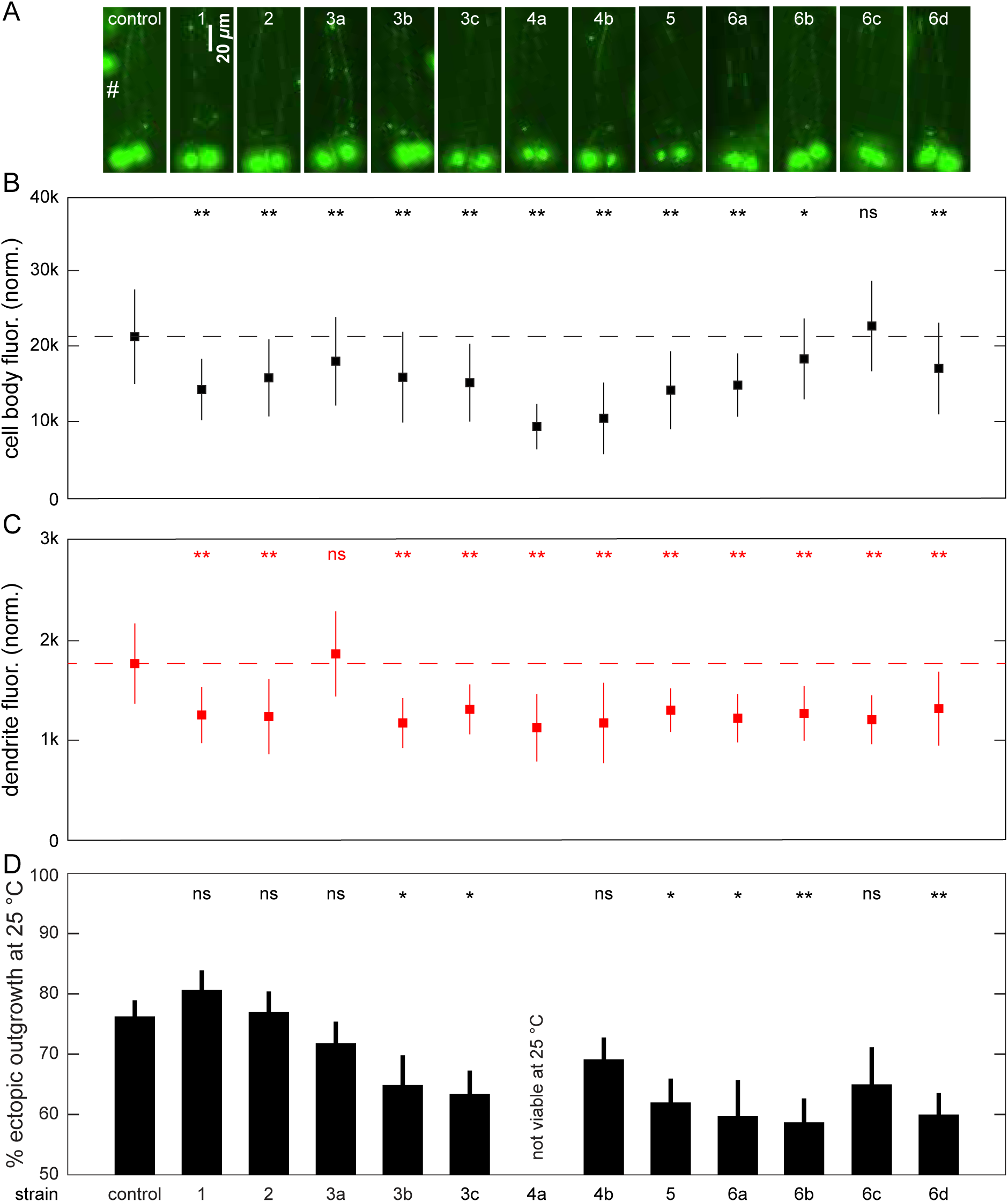
Fluorescence proxy enables isolation of mutants with reduced conditioned regenerative potential. (a) ASJ images in control and post-mutagenesis animals. Fluorescent bead indicated by #. Identical brightness, contrast for all images. (b) Quantification of cell body fluorescence in part A. Reduced fluorescence in 11 mutant lines. (c) Quantification of dendrite fluorescence in part A. Reduced fluorescence in 11 mutant lines. (d) Ectopic outgrowth frequency for control and post-mutagenesis strains. Reduced ectopic outgrowth in 6 lines. Fluorescence measurements: *n =* 50. Ectopic outgrowth: *60* ≤ *n ≤ 94* for strains 3b, 6a, 6c; *n ≥ 150* for other strains. Strain 4a not viable at cultivation temperature. Data represented as average ± SD. * *p < 0.05, ** p < 0.001*. Control (*tax-2*) images and fluorescence data from Fig. 1.

Mutations to genes involved in the sensory pathways (*i.e.*, conditioning mutations) decrease ASJ neuronal activity and trigger regeneration. These sensory mutations, including the mutations in Fig. 1, also alter axon morphology in *C. elegans* sensory neurons and produce ectopic outgrowths (Coburn and Bargmann, 1996; Peckol, *et al*., 1999). We found that ectopic outgrowth extensively shares genetic and molecular mechanisms of axon growth with conditioned regeneration (Chung, *et al*., 2016). Thus, ectopic outgrowth indicates conditioned regenerative potential. As shown in Fig. 4d, six of the strains we isolated exhibit significantly reduced frequencies of ectopic outgrowth compared to the original pre-mutagenized strain (*p < 0.05* or *0.001*), suggesting that these strains possess a mutation that restrains conditioned regeneration in the ASJ. As we previously found, fluorescence is not always an accurate predictor of conditioned outgrowth. We identified some strains with significantly reduced cell body and dendrite fluorescence but unchanged ectopic outgrowth frequency. We expect these strains to carry a mutation that decreases *trx-1* expression without impact to the conditioning pathway. In summary, our results demonstrate the utility and limitations of our fluorescence proxy as an indicator of ASJ intrinsic regenerative capacity.

## Discussion

The change in expression of TXN following neuronal injury has been widely characterized in mammalian models. TXN induction occurs after a variety of insults, including ischemia (Tomimoto, *et al*., 1993) and oxidative stress (Sugino, *et al*., 1999). Importantly, motor and sensory nerve axotomy in rats leads to significantly increased levels of TXN (Mansur, *et al*., 1998; Stemme, *et al*., 1985). Likewise, we note that laser surgery of the ASJ neurites upregulates *trx-1* expression, as indicated by *trx-1::gfp* reporter. Our study offers three lines of evidence for associating this enhanced *trx-1* expression with the conditioned state. First, changes due to conditioning, including upregulation of RAGs, preferentially occurs after lesioning the sensory neurite rather than the central, or synaptic, neurite. Severing the sensory dendrite strongly upregulates *trx-1* in nearly all ASJ neurons while severing the axon upregulates expression in only 50% of neurons to a lesser degree (Fig. 2b). Second, each mutation we tested that improves regeneration (Chung, *et al*., 2016) by conditioning the ASJ neuron also upregulates expression of *trx-1* as measured by GFP (Fig. 1b) and FISH (Figs. 1cd). Third, *trx-1* expression generally corresponds with conditioned regenerative potential (Fig. 1b). Out of the mutations tested, the *tax-2* and *tax-4* mutations brighten the ASJ the most and produce the highest levels of ectopic outgrowth and regeneration (Chung, *et al*., 2016). However, this relationship is not comprehensive (Fig. S1). The *tax-2* and *tax-4* mutations produce marginally better conditioned regeneration compared to other mutations, and the *unc-36* mutation increases *trx-1* expression but minimally triggers regeneration (Chung, *et al*., 2016). This differential alteration of *trx-1* expression and regeneration suggests a divergence of pathways underlying *trx-1* expression and conditioning.

To our knowledge, the role of *trx-1* in neuronal regeneration has not directly been studied, although prior evidence suggests its role as a mediator. Overexpressing TXN in mice can promote neuroprotective effects following stroke (Takagi, *et al*., 1999), and TXN is required for nerve growth factor enhancement of nerve outgrowth in a tumor cell line (Bai, *et al*., 2003). We directly test the role of *trx-1* in modulating conventional and conditioned regeneration in the ASJ (Fig. 3). We show that *trx-1* inhibits conventional regeneration, consistent with findings in the PLM sensory neuron in *C. elegans*. Animals defective in thioredoxin reductase *trxr-1*, which enables *trx-1* activity (Stenvall, *et al*., 2011), display increased PLM regeneration (Kim, *et al*., 2018). We also show that *trx-1* mediates *dlk*-independent conditioned regeneration, consistent with the upregulation of *trx-1* expression following conditioning interventions. Lesion conditioning is enabled by changes in transcription that lead to an enhancement of regeneration (Hoffman, 2010). The mammalian TXN gene contains a regulatory region with cAMP responsive element (CRE) through which nerve growth factor acts to induce nerve outgrowth (Bai, *et al*., 2003). The *trx-1* gene has a similar consensus CRE sequence, CGTCA (Montminy, *et al*., 1986; Park, *et al*., 2021). We speculate CRE-binding protein (CREB), an integral transcription factor for many biological processes, could regulate *trx-1* transcription through this region to enact the conditioning response. The cAMP pathway is one of the best-described mechanisms for enhancing conditioned regeneration (Neumann, *et al*., 2002; Qiu, *et al*., 2002), and in *C. elegans*, cAMP signaling triggers conditioned regeneration (Chung, *et al*., 2016). Likewise, we recently found that CREB enhances conditioned regeneration (Wang, *et al*., 2023). Thus, cAMP and CREB may contribute to conditioned regeneration in part via *trx-1* activation through its CRE region.

The mechanisms by which *trx-1* differentially affects conventional and conditioned regeneration requires additional study. Dendrite injury preferentially upregulates *trx-1*, which would presumably enhance conditioned regeneration over conventional regeneration. This matches the perturbation and phenotype of a conditioning lesion. The conventional and conditioned regeneration pathways are fundamentally neurite growth mechanisms, and they likely converge or regulate each other to produce this growth. Their interaction may favor one pathway over the other depending on context, such as neuron development (regeneration recapitulates some aspects of development), regeneration, or even cell death and stress response. Both *trx-1* and *dlk-1* are known to interact with JNK/MAPK signaling cascades that initiate cell death in response to stress (Fujino, *et al*., 2007; Nix, *et al*., 2011). The specific interaction of *dlk-1* and *trx-1* to drive their biological functions remains undefined. However, our results in Fig. 3 suggest that the inhibition of conventional regeneration by *trx-1* acts upstream or converges to *dlk-1* given that loss of *trx-1* only significantly increases conventional regeneration with functional DLK.

Redox activities of thioredoxin in *C. elegans* are important for modulating biological processes (Sanzo-Machuca, *et al*., 2019). In mammalian models, the redox-related pathways of TXN have been heavily studied, particularly in regulating apoptosis and protecting against neurodegenerative disorders. It is not surprising that *trx-1* plays an integral biological role in these processes since a hallmark of cell death and neurodegeneration is an accumulation of damaged proteins, reactive oxygen species, and oxidative stress (Lin and Beal, 2006; Mattson and Magnus, 2006). Thioredoxin inhibits apoptosis signal-regulating kinase 1 (ASK1) to mediate endoplasmic reticulum stress-induced apoptosis (Nishitoh, *et al*., 2002; Saitoh, *et al*., 1998) and can delay retinal neurodegeneration in the Tubby mouse model (Kong, *et al*., 2007; Kong, *et al*., 2010). It is still unknown how these redox functions dictate regenerative potential, but stress response and cell death pathways are important modulators of regeneration (Pinan-Lucarre, *et al*., 2012). We speculate that *trx-1* may impinge on or regulate these pathways to affect axon regeneration.

Using our fluorescence proxy, we isolated twelve mutant lines with significantly decreased ASJ fluorescence. Six of those lines display significantly reduced ectopic outgrowth, which is related to conditioned regeneration, indicating that they contain a mutation in the conditioning pathways. Additionally, we developed a fluorescent bead normalization protocol to facilitate quantitative comparison of fluorescence in genetic backgrounds or under different interventions. These tools enable us to pursue a rapid, yet quantitative approach to the question of neuronal regeneration. Our approach is complimentary to established techniques for identifying RAGs, such as microarray analysis (Costigan, *et al*., 2002). In contrast to these RAGs, genes identified by our approach are highly likely to functionally underlie regeneration, by virtue of their involvement in ectopic outgrowth.

Our proxy demonstrates significant potential as we isolated several strains with reduced outgrowth potential by selecting for reduced ASJ fluorescence. However, we did identify equally as many dim strains without reduced outgrowth potential (Fig. 4d). These findings are not unexpected since we introduce stochastic mutations throughout the genome and screen with a fluorescent indicator that only roughly corresponds with regenerative capacity. False positives can be a common drawback to forward genetic screens under mutagenesis due to their stochastic nature. In these isolated lines, we expect that EMS disrupted pathways directly related to *trx-1* expression, genes that interact upstream of *trx-1*, or the transgenic label itself, thus causing a significant reduction in fluorescence. These effects can be mitigated by pursuing a reverse genetic screen under RNAi to test genes that have an expected role in conditioned regeneration. A similar screen was successfully carried out on transcription factors regulating non-conditioned *trx-1* expression (Gonzalez-Barrios, *et al*., 2015).

Additionally, we screen for decreased fluorescence by eye under the current workflow. This reduction in fluorescence is often difficult to distinguish by eye. Also, the range of fluorescence of the starting strain and mutant often overlap, which prevented outcrossing and identification of the causative mutations. We could improve the throughput and probability of success by implementing real-time quantitative measurements to more precisely screen for changes in fluorescence. For instance, we could utilize microfluidic devices to rapidly screen through animals for a reduction in fluorescence or implement whole plate imaging (Wang, *et al*., 2023) to quickly image entire populations of animals to select for decreased fluorescence.

### Outlook

Studying regeneration *in vivo* remains difficult for experimental reasons and due to interacting pathways yet to be explored and fully defined. We have developed a single-cell model to study the conventional and conditioned regeneration pathways in the *C. elegans* ASJ neuron and a proxy to identify genes and potential therapeutic targets involved. Our study could be extended in multiple directions. First, the mechanisms through which *trx-1* (and its redox activities) modulates regeneration is of particular interest because *trx-1* specifically enhances conditioned, but inhibits conventional, regeneration. Second, the mechanisms that underlie *trx-1* expression and their relation to conditioning pathways should be clarified. Third, while *trx-1* expression is specific to the ASJ, our approach could be extended to other neurons by labeling with fluorescent reporters driven by their RAGs. Neuron types and even subtypes have distinct regeneration capacities (Duan, *et al*., 2015), but broad regeneration studies in multiple neurons are lacking. Examining regeneration in other neurons may illuminate new regeneration pathways. Our work thus represents an important first step in identifying undiscovered modulators of neuron regeneration and in establishing new approaches for rapidly studying regeneration.

## Abbreviations

CNS: central nervous system
EMS: ethyl methanesulfonate
DLK: dual-leucine kinase
TXN: thioredoxin
cAMP: cyclic adenosine monophosphate
CRE: cAMP responsive element
CRE-binding protein: (CREB)
RAG: regeneration associated gene
RNAi: ribonucleic acid interference
a+d: concomitant axon + dendrite surgery
FISH: fluorescence in situ hybridization

## Acknowledgements

We thank members of the Chung laboratory for taking part in the revision process. We thank Leilani Schulting for assistance with data postprocessing. We acknowledge Peter Swoboda (Karolinska Institute, Huddinge, Sweden) for providing OE3417 and OE3265. We would like to also thank Antonio Miranda-Vizuete (Biomedicine Institute of Sevilla, Seville, Spain) for providing VZ763, VZ748, and invaluable feedback on the manuscript.

