## Supplementary figures and images for "Expression of thioredoxin-1 in the ASJ neuron corresponds with and enhances intrinsic regenerative capacity under lesion conditioning in *C. elegans*"

### Supplemental Figure 1

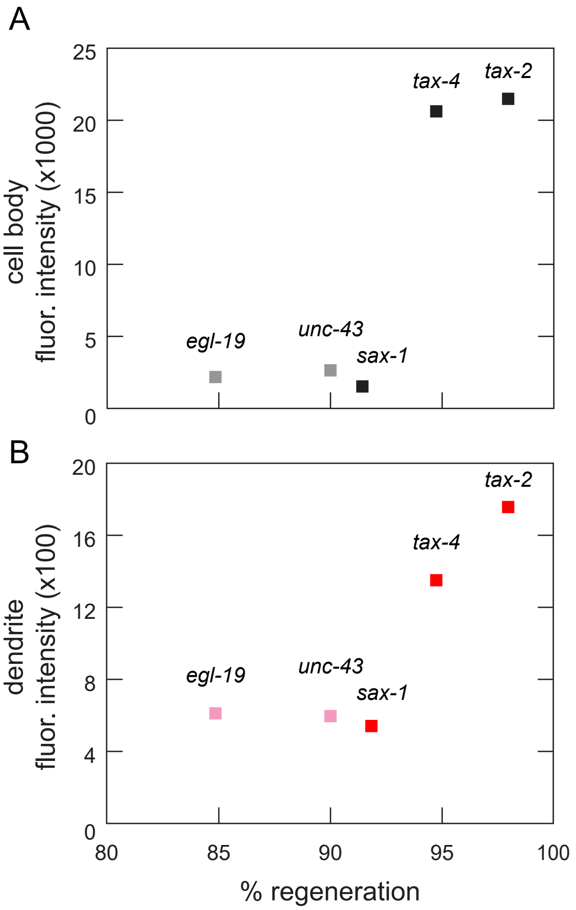


Figure S1
*Grooms, et al*
