## Supplemental Table 1 and Supplemental Figure 1 caption for "Expression of thioredoxin-1 in the ASJ neuron corresponds with and enhances intrinsic regenerative capacity under lesion conditioning in *C. elegans*"

| Probe # | Probe sequence |
| --- | --- |
| 1 | gttaccatatcagcaagctc |
| 2 | agttgcatcgtttcaacatt |
| 3 | attgctcaaagtcactctga |
| 4 | aatgatcttctccggatgtt |
| 5 | cgcaccaagttgcatagaaa |
| 6 | aatggtgcaattgctttgca |
| 7 | cctttgtgagttgtagctaa |
| 8 | atcgacatcaactttgcaga |
| 9 | tggaacaaagatcttccgct |
| 10 | aagtcggcatcatcttgaca |
| 11 | cgtctccattcttggtgaaa |
| 12 | acactttttgacgcagttcg |
| 13 | cattgagcagatacgtgctc |
| 14 | ggtttaaaagcagatggtcg |
| 15 | cggaagagctcaatgagcag |
| 16 | gagcggggtttaattgtagg |

**Table S1:** List of generated FISH probes used to target *trx-1*.

**Figure S1: Expression of *trx-1* is positively associated with regenerative potential.** Sensory mutations enhance ASJ regeneration and *trx-1* expression in cell body (a) and dendrite (b). Red and black icons denote ASJ is labeled with *ofIs1*; gray and pink, *ofIs4*.
